# CAR-T cells locally delivered in porcine decellularized matrix hydrogels enhance survival in post-resection glioblastoma

**DOI:** 10.1101/2025.02.11.637648

**Authors:** Meghan T. Logun, Sabrina L. Begley, Kelly Hicks, Jungmin Park, Logan Zhang, Zev A. Binder, Donald M. O’Rourke

## Abstract

Patients diagnosed with glioblastoma (GBM) receive a devastating prognosis of less than 15 months, and recurrence of GBM is most often local, suggesting that regional therapies would serve both immediate and long-term needs of patients. Here, we investigate a biomaterials-based approach for local delivery of chimeric antigen receptor (CAR) T cells using a murine model of partial GBM resection that mimics patient recurrence. We demonstrate that hydrogel delivery of CAR T cells directly into the intracranial resection cavity can stably implant cellular immunotherapies against CNS solid tumors, and significantly prolongs survival in recurrent GBM-bearing mice compared to those receiving resection alone.

**One Sentence Summary:** During resection surgeries for brain tumors, delivering cell therapy at time of surgery could improve outcomes for patients.

## INTRODUCTION

Patient prognosis for glioblastoma (GBM), the most common and lethal of malignant primary brain tumors, is extremely poor. Median survival for newly diagnosed GBM remains less than two years(*1*). No current treatments are curative and standard-of-care (SOC) regimens including resection and chemoradiation are often temporizing measures due to the diffusely infiltrative nature of the disease. Only three new treatments have been FDA-approved for GBM since 2005: temozolomide (TMZ), bevacizumab, and tumor-treating fields(*1–4*), of which only TMZ and tumor-treated fields demonstrate prolongation of survival. Characterization of GBM cells has resulted in the identification of several targets amenable to molecular targeting using immunotherapies such as chimeric antigen receptor (CAR) T cells. CAR T cells have demonstrated impressive patient responses with hematological malignancies. Unfortunately, the efficacy of CAR T cells against GBM have been limited thus far due to the immunosuppressive tumor microenvironment (TME), adaptive resistance from the tumor, inadequate trafficking of CAR T cells from peripheral administration, limited diffusion across the blood-brain- and blood-tumor-barriers, and lack of CAR T cell persistence(*5*).

Our group performed the first-in-human trial (NCT02209376) of autologous T cells redirected to EGFR variant III for recurrent GBM, finding that intravenous (IV) infusion resulted in on-target activity in the brain. However, the CAR T cells ultimately could not overcome the adaptive changes in the local TME and led to only limited clinical efficacy(*6*). Another landmark trial demonstrated that local delivery of interleukin 13 receptor subunit alpha (IL13Rα2)-targeted CAR T cells was associated with elevated inflammatory cytokine levels and increased presence of CAR+ T cells in cerebrospinal fluid, but disease progression was only delayed and not prevented(*7*). Recurrence is currently inevitable despite SOC approaches, but novel local delivery methods could enhance cellular immunotherapy efficacy. Delivering CAR T cells via a biomaterial carrier implanted into the post-resection cavity would not only overcome the inadequate trafficking from peripheral administration but also enhance penetration into the cavity by bypassing the heterogeneous blood-tumor- and blood-brain-barriers.

Hydrogels were classically used as drug delivery systems for hydrophilic drugs. However, advancements in cell culture efforts have developed cell carrier systems that are widely used in stem cell treatments for major injuries and defects(*8*). Hydrogels with varying components, including collagens, hyaluronic acid (HA), chitosan, and decellularized extracellular matrices (dECMs), have been applied to nerve cell regeneration, traumatic brain injury (TBI), and spinal cord repair(*9, 10*). Decellularization of tissues provides ECM scaffolds for regeneration: eliminating cellular and genetic components while preserving the composition, structure, and function of native ECM proteins(*11*). Recent technological advancements have enabled the decellularization of large organs such as liver, lung, kidney, and heart. These decellularized scaffolds have been used for applications including wound healing, bone defects, nerve injuries, heart disease, and whole-organ tissue regeneration(*12–16*).

Uniquely, the ECM of brain tissue contains fewer fibrous structural proteins, such as collagen or elastin, and more proteoglycans compared to other tissues(*17*). Proteoglycans and their glycosaminoglycans (GAG) side chains profoundly influence neuronal cell behavior via adhesion and neurite outgrowth(*17*). Due to these specialized features, dECM is gaining popularity in preclinical models of neural injury and repair. dECM processing largely preserves collagen fibrils and glycoproteins - the components responsible for the reassembly into a hydrogel upon injection *in vivo*(*18–21*) – while retaining physiologically relevant amounts of proteoglycans, ECM regulators, and ECM-associated proteins that other constructs (such as the non-neuronal Matrigel (Corning)) lack(*22*). Current ECM applications include the intraoperative use of fibrin sealants, inspired by the natural clotting factors in the bloodstream, to increase post-resection hemostasis in the operative cavity. This delivery of therapeutically relevant scaffolding into the resection cavity could provide a helpful microenvironment for implanted cell therapeutics to thrive.

In this study, we fabricated a practical dECM hydrogel platform for the locoregional delivery of CAR T cells into a mouse model of post-resection GBM. These dECM biogels provided a physiologically relevant scaffold for the implantation and migration of CAR T cells into surgical margins, enhancing antitumor responses and survival outcomes in dECM-treated groups compared to peripherally administered CAR T cells. Furthermore, local dECM hydrogel delivery allowed for equivalent efficacy outcomes at lower doses than required for peripheral administration.

## RESULTS

### Development and validation of dECM platform

GBM recurrence is inevitable even with the best available SOC treatments that include at least one, if not multiple, surgical resections. As shown in Fig. 1A, resection of an orthotopic, intracranial GBM in mice established a model for post-resection and recurrent GBM. Cells left within the resection margins repopulated and lead to recurrence within 30d. To test local delivery using the resection surgery as an opportunity for early treatment, we encapsulated EGFRvIII targeting CAR T cells into decellularized brain matrix hydrogels to administer directly into th tumor resection cavity (Fig. 1B).

**Fig. 1.**
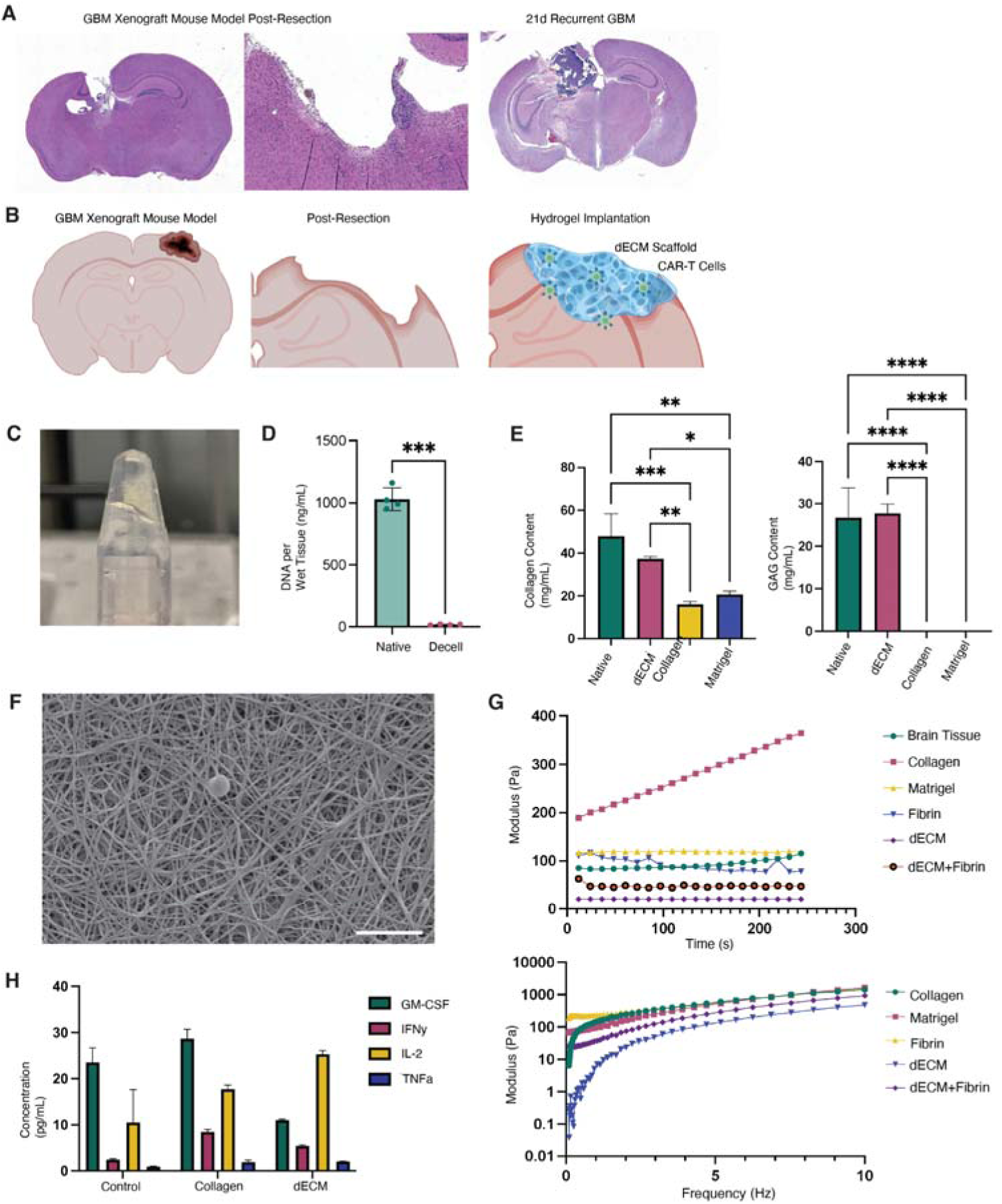
Characterization of GBM recurrence model and design of decellularized brain extracellular matrix hydrogels (dECM). (**A**) Hematoxylin and eosin (H&E) staining of intracranial GBM xenograft tumor cavity directly after surgical resection procedure (left). Middle inset shows a magnified image of subtotal surgical resection with GBM cells remaining in margins. Tumors demonstrate recurrence within 21d of resection procedure (right). (**B**) Schematic illustration of local delivery of CAR T cells directly into the resection cavity after surgery for delaying GBM recurrence and extending survival outcomes. (**C**) Photo of decellularized hydrogel solution maintaining shape after gelation procedures. (**D**) DNA quantification in native and decellularized porcine brain tissues. Statistical significance determined by Student’s t-test, \*\*\**p* < 0.001. (**E**) Quantification of ECM components collagen (left) and glycosaminoglycan (“GAG,” right) in native tissue, decellularized tissue, or biosimilar hydrogel controls EZ-Coll and Matrigel. Statistical significance determined by one-way ANOVA, \**p* < 0.05; \*\**p* < 0.01; \*\*\**p* < 0.001; \*\*\*\**p* < 0.0001. (**F**) Scanning electron micrograph of dECM hydrogels showing network of fiber structures. Scale bar 50 µm. (**G**) Rheology trace of storage modulus over time (top) and frequency (bottom) demonstrating dECM hydrogel gelation stiffness and shear properties compared to native brain tissue and additional biosimilar hydrogel networks. (**H**) Quantification of inflammatory cytokines GM-CSF, IFNγ, IL-2, or TNFα released by cells after encapsulation in EZ-Coll, dECM, or no gels demonstrating a lack of significant inflammation brought on by encapsulation. Statistical significance determined by one-way ANOVA and ‘ns’ indicates no significance.

The decellularization and solubilization of the porcine brain (Fig. S1) created a viscous, pre-gel solution that formed a self-supporting soft hydrogel after crosslinking at 37°C for 5 minutes (Fig. 1C). Decellularization was confirmed through a lack of DNA detected in the pre-gel solution as compared to source native brain material (Fig. 1D). As decellularized matrices are rarely homogeneous in composition, protein composition was quantified across batches using colorimetric assays, finding mostly collagens and glycosaminoglycans (GAGs) responsible for the hydrogel backbone (Fig. 1E). Our dECM hydrogels were compared to Matrigel, EZ Coll (Advanced Biomatrix), and fibrin (Sigma Aldrich) alone on the basis of protein composition, stiffness and cytokine release. To increase cohesion and achieve hemostasis in a normally bloody resection cavity, fibrinogen was added at physiologic levels. This was planned to improve molding and adhesion of the dECM to the walls of an irregularly shaped resection cavity while preventing dilution of delivered therapies. The addition of fibrin to strengthen adhesion in the resection cavity has been evaluated separately and showed no influence on T cell viability(*23–25*).

To ensure the appropriate structure and function of our decellularized brain extracellular matrix (dECM) as a T cell transplantation scaffold, we performed scanning electron microscopy (SEM), rheological analysis, and cytokine release quantifications. The solidified hydrogel network can be visualized using SEM with pore sizes ranging from 5 to 30 µm, which are large enough for T cells to move through (Fig. 1F). Storage modulus (G’) was measured over time to ensure that gels maintained viscoelastic solid behavior and had a stiffness profile comparable to native brain tissue (Fig. 1G). Brain tissue is notoriously soft, with compressive moduli ranging around 0.5 – 5 kPa, and the porcine dECM devoid of cellular material fell at the lower end of that range in our hands(*23–25*). A second rheological test was performed, measuring the storage modulus (G’) in a frequency sweep test to observe the viscoelastic spectrum of each gel system under oscillatory stress (Fig. 1G). The addition of fibrin to the dECM generated a storage modulus closer that of native brain and a solution that is more elastic but less viscous than the dECM alone. Therefore, this combination was used for all subsequent in vivo studies. Finally, untransduced T cells were encapsulated into dECM hydrogels for 24h and then media was analyzed for release of inflammatory cytokines. Compared to control unencapsulated T cells or T cells left in collagen-only hydrogels, dECM-encapsulated T cells had a lower inflammatory cytokine presence and greater IL-2 enrichment (Fig. 1H).

### Released CAR T cells maintain viability and cytotoxic capability

EGFRvIII CAR T cells were used to test the ability of dECM hydrogels to maintain T cell viability and release *in vitro* utilizing Transwell assays. CAR T cells were first expanded and transduced with a lentiviral vector containing a chimeric scFv targeting the EGFR epitope 806(*26*) and then EGFR CAR+ populations were analyzed using flow cytometry (Supp. Fig. 2). When 1.5M T cells were incubated in each treatment paradigm within the Transwell inserts, all gels had similar release kinetics compared to the unencapsulated T cells moving through the Transwell membranes (Fig. 2A). Cytocompatibility of dECM hydrogels was compared to other commercially available cell matrix formulations using colorimetric cell viability assays after encapsulation. CAR T cells retained high cell viability over 96h in all hydrogel treatments as compared to unencapsulated gels (Fig. 2B). Released cells were collected from Transwell assays after 48h and placed into tumor killing assays using an impedance-based platform. Cytotoxic ability was not significantly impacted by 48h encapsulation across hydrogel matrices or when compared to unencapsulated CAR-T cells at 1:1 E:T ratio against U87MG tumor cells (Fig. 2C).

**Fig. 2.**
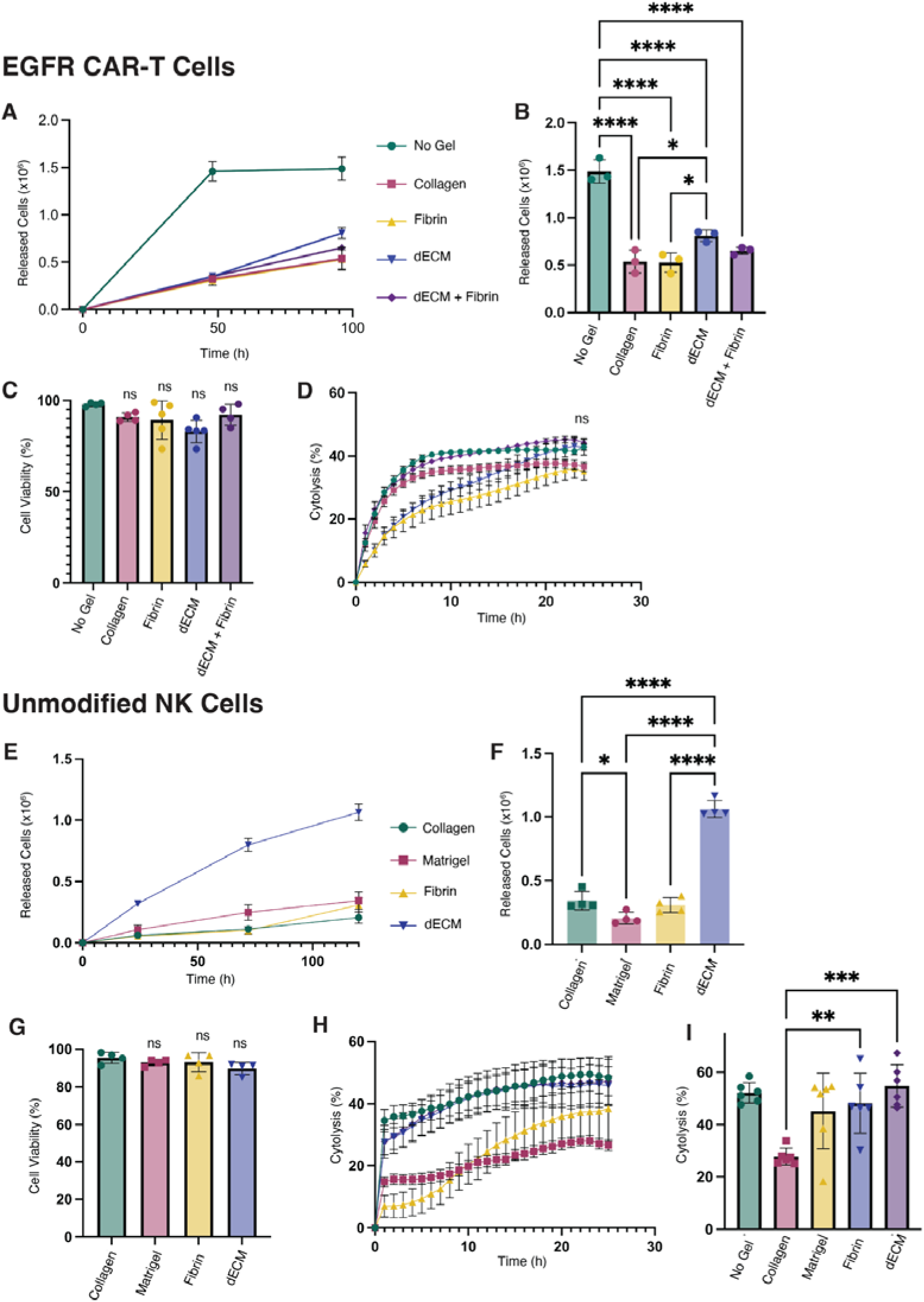
dECM hydrogel encapsulation allows for reliable bulk release of immunotherapeutic cells compared to biosimilar control hydrogels. (**A**) 2.0×10^6^ EGFRCAR-T cells were encapsulated within hydrogels in Transwell Assays to observe cell release kinetics over time in culture and at (**B**) assay endpoint of 96h. Statistical significance was calculated using one-way ANOVA, \**p* < 0.05; \*\*\*\**p* < 0.0001. (**C**) Cell viability of released cells was quantified using live/dead staining and then cells released after 48h were placed into (**D**) impedance-based tumor killing assays at 1:10 E:T against target U87vIII cells. Statistical significance determined by one-way ANOVA and ‘ns’ indicates no significance. (**E**) To observe any effects of cargo on release kinetics, normal donor NK cells were encapsulated into hydrogel treatments and monitored for release over time and (**F**) at 120h endpoint. Statistical significance was calculated using one-way ANOVA, \**p* < 0.05; \*\*\*\**p* < 0.0001. (**G**) NK cell viability was quantified in released cells and released viable cells were seeded into impedance tumor killing assays at 1:10 E:T against target U87vIII cells (**H-I**). Statistical significance was calculated using one-way ANOVA, \*\**p* < 0.01; \*\*\**p* < 0.001; ‘ns’ indicates no significance.

To demonstrate the utility of dECM hydrogels as cell carrier options for other cell therapy platforms, we analyzed the viability, release, and cytotoxicity of NK cells encapsulated in various hydrogel carriers. Interestingly, dECM hydrogels enabled a significantly greater release of NK cells over time when compared to commercially available competitor hydrogels, although all released cells were equally viable across groups (Fig. 2D-E). In killing assays conducted 24h after release, dECM-released NK cells significantly outperformed cells from other matrices and endpoint cytolysis values were similar to cells that had not experienced gel encapsulation (Fig. 2F-G). The observed differences in NK release from hydrogel matrices of varying compositions align with prior observations that immune cells are sensitive to changes within the microenvironment and as such, this should be considered in the design of delivery vehicles.

### Establishment of intracranial resection surgeries and recurrence models in mice

Before evaluating antitumor efficacy, we developed an NSG murine model of intracranial recurrent GBM where survival was not significantly enhanced by surgery alone (Supp Fig. 3). Tumors were generated via intracranial injection, and after seven days of growth, suction ablation was used to resect a 1.5mm x 2mm section of tissue from where tumor was observed on bioluminescence imaging (BLI). This created a subtotal resection cavity with tumor cells left in the margins, mimicking human GBM resection results (Fig 1A). Tumors demonstrated significant regrowth within 7d of resection, and continued to progress until a humane endpoint was reached. Post-resection survival was minimally extended compared to no treatment at all, mimicking human GBM clinical paradigms. Pathologic examination of endpoint tumors between control untreated and surgery treated animals were indistinguishable from each other in tumor bulk (Supp Fig. 3). To ensure dECM hydrogels containing EGFR CARs could be safely implanted, empty dECM hydrogels were examined in both flank and intracranial injections in order to observe the effects on local healthy tissue (Supp. Fig. 4). No local inflammation was observed in flank injections after 7d, and mice that received intracavitary dECM only after intracranial resections exhibited tumor recurrence at the same rate as resection only mice (Supp. Fig. 4), indicating that empty hydrogels had no effect on recurrence dynamics in these models.

**Fig. 3.**
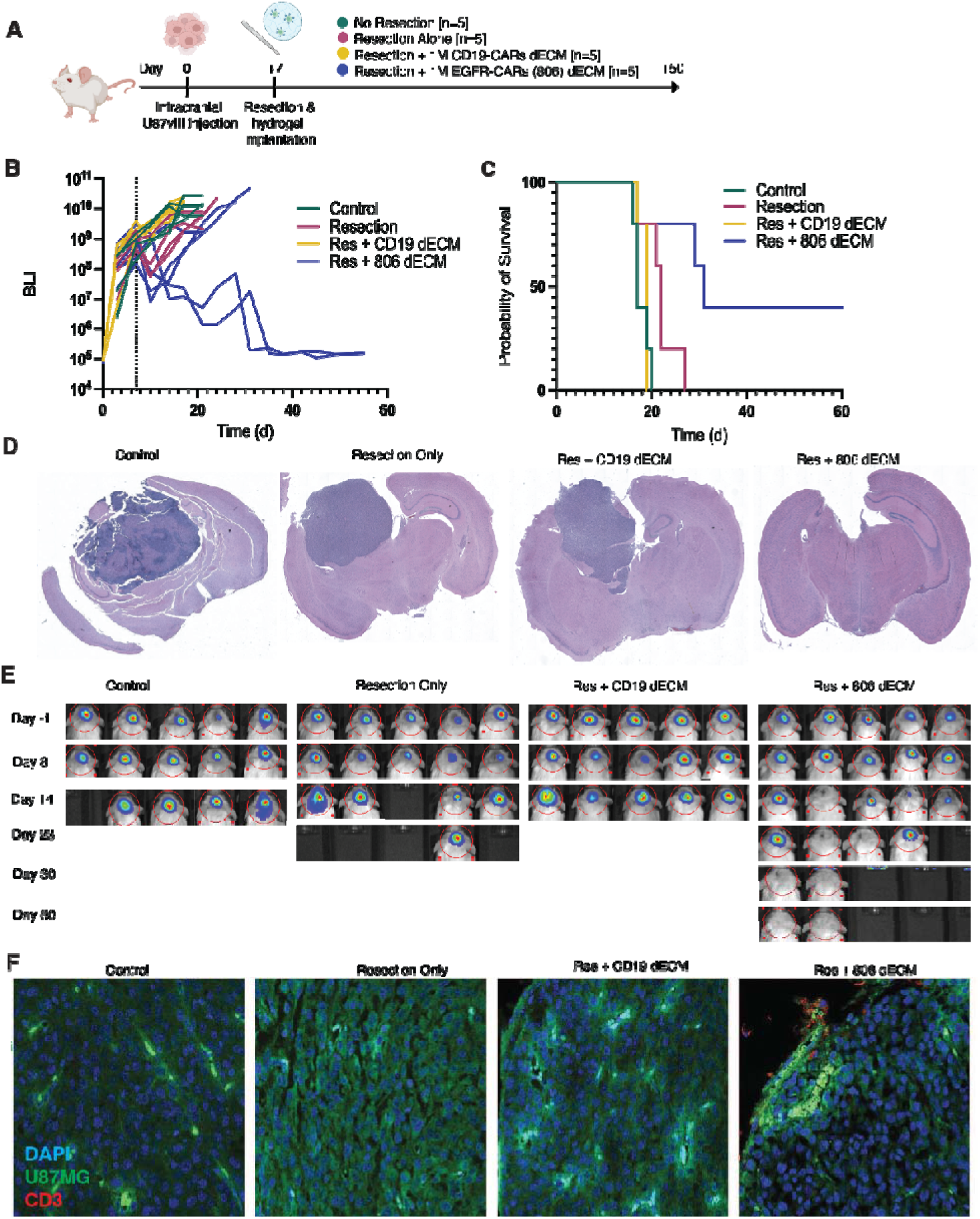
Locoregional delivery of on-target EGFR-CAR T cells enhances survival in post-resection xenograft mice. (**A**) Schematic representation of the experimental design. U87vIII cells were transplanted into NSG mice. After tumor growth over 7d, mice were given no additional treatment, incomplete resection surgery alone, resection surgery with locoregional dECM hydrogels containing 1×10^6^ off-target CD19-CAR T cells, or resection surgery with locoregional dECM hydrogels containing 1×10^6^ on-target EGFR-CAR-T cells (n=5 per group). Mice were followed with BLI until 60d or until humane endpoints were reached. (**B**) BLI signal representing tumor burden measured in individual mice over time. The vertical line indicates time of surgical resection and dECM treatment. (**C**) Kaplan-Meier survival curves of control, resection only, resection with CD19 dECM or resection with EGFR dECM-treated mice (n=5 per group). Data are analyzed with the log-rank Mantel-Cox test. (**D**) Representative images of BLI of all mice over time, red circles highlight tumor location. (**E**) Representative H&E images of endpoint tumors acquired from each treatment group. (**F**) Representative images of tumor containing tissue with immunohistochemical staining for CD3+ cells (red). U87vIII tumor cells express GFP (green).

### In vivo efficacy of locoregionally delivered EGFRvIII CAR-T cells

*In vivo* antitumor efficacy was evaluated using a U87MG-EGFRvIII+ xenogeneic orthotopic model in NSG mice. Seven days after inoculation with U87MG-EGFRvIII+ cells, mice were randomized to an experimental group (no treatment, resection alone, resection + 1×10^6^ CD19-CAR T cells in dECM, or resection + 1×10^6^ EGFR-CAR T cells in dECM) and treated accordingly (Fig. 3A). In the groups that received a resection procedure, mice either received a resection alone or a resection followed by an intracavitary injection of dECM loaded with either on-target EGFR-CARs or negative control CD19 CAR T cells. Cell surface staining of the transduced CAR T cells confirmed expression of the CAR construct in 40-60% of cells (Supp Fig. 2B). Mice were followed for a maximum of 60 days after tumor inoculation and tumor growth or recurrence was monitored via BLI signals from CBG+ U87MG-EGFRvIII+ cells. Locally delivered on-target EGFR-CAR T cells delivered via dECM hydrogel significantly enhanced survival outcomes compared to both dECM-loaded off-target CD19-CARs and resection alone (Fig. 3C). Positive responders to on-target local therapy experienced durable responses as seen on both BLI and in their healthy weight conditions over the 60 days of treatment (Fig. 3B-D). Brain tumor tissues were harvested at study endpoints, and tumor-bearing brain slices with hematoxylin and eosin (H&E) stains were imaged to examine tumor morphologies across treatment groups (Fig. 3E). In tumor-bearing brain sections stained for CD3+ cells, residual CD3+ T cells were only detected in endpoint animals that were treated with EGFR-CARs loaded into dECM hydrogels (Fig. 3F).

### Intracavitary injection of CAR-T loaded hydrogels compared to intravenous treatment

The efficacy of locally delivered CAR T cells was compared against peripherally delivered CAR T cells at the same dose using our previously described resection xenograft model, Seven days after inoculation with U87MG-EGFRvIII cells, animals received partial resection procedures (Fig. 4A). In groups that underwent resection, mice either received a resection alone, a resection with intracavitary injection of dECM loaded with EGFR-CAR T cellss, or a resection with a peripheral injection of EGFR-CAR T cells delivered via tail vein. Animals were followed for a maximum of 60 days after tumor inoculation and tumor recurrence was monitored via BLI signals from CBG+ U87MG-EGFRvIII+ cells (Fig. 4B-D). After the local delivery of CAR T cells, a reduction in BLI signals was seen within days and persisted until the study endpoint (Fig. 4B, 4D). Compared to peripherally delivered CAR T cells at the same dose, the locally delivered CAR T cells significantly increased survival probability (Fig. 4C). Brain tumor tissues were harvested at study endpoint, and tumor-bearing brain slices with H&E stains were imaged to examine tumor areas across treatment (Fig. 4E). Tumor-bearing tissues were also stained for the presence of CD3+ cells, which were only detected in animals treated with EGFR-CAR T cells in dECM hydrogels (Fig. 4F).

**Fig. 4.**
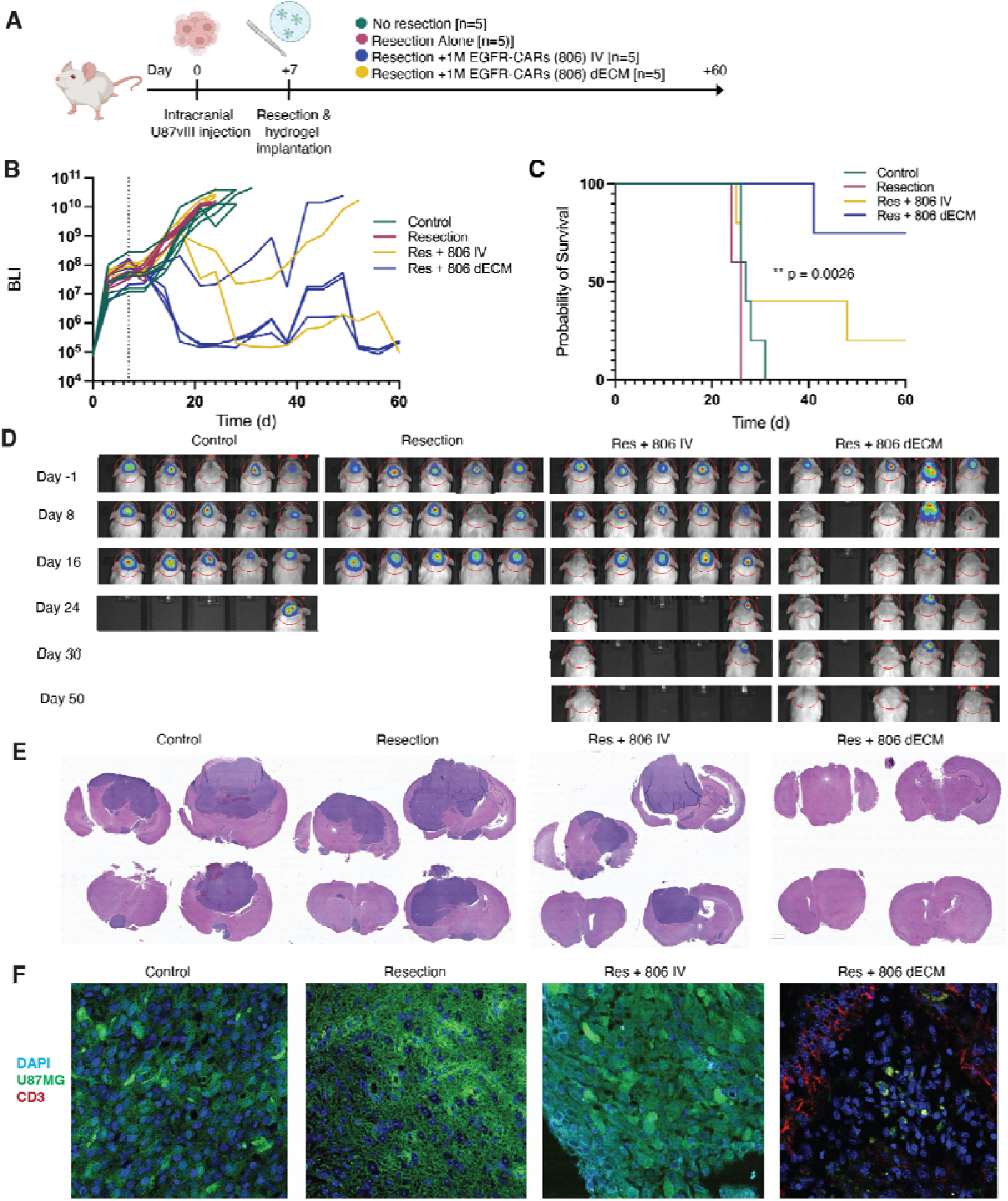
A singular locoregional dose of EGFR-CAR-loaded dECM hydrogels controls recurrence in post-resection GBM xenograft mice compared to equivalent dose administered through clinical standard intravenous delivery. (**A**) Schematic representation of the experimental design. U87vIII cells were transplanted into NSG mice. After tumor growth over 7d, mice received either no treatment, incomplete surgical resection alone, surgical resection with intravenous 1×10^6^ EGFR-CAR T cells, or surgical resection with locally injected dECM hydrogels containing 1×10^6^ EGFR-CAR T cells (n=5 per group). Mice were followed with BLI until 60d or until humane endpoints were reached. (**B**) BLI signal representing tumor burden measured in individual mice over time. Arrow indicates time of surgical resection and treatment. (**C**) Kaplan-Meier survival curves of control, resection only, resection with IV EGFR-CARs, or resection with EGFRgel treatment. Data are analyzed with the log-rank Mantel-Cox test. (**D**) Representative images of BLI of all mice over time, red circles highlight tumor location. (**E**) Representative H&E images of endpoint tumors acquired from each treatment group. (**F**) Representative images of tumor containing tissue with immunohistochemical staining for CD3+ cells (red). U87vIII tumor cells express GFP (green).

### Local delivery enables efficacy at lower dosing

Within the field of cell therapies, recommended dosing remains an effervescent question. The answer varies greatly depending on the type of therapy, patient disease criteria, and the route of administration. In order to evaluate how the route of administration, and therefore delivery vehicle, impacts the necessary dose requirement, we prepared a third cohort of xenografted mice with partially resected GBM. NSG mice with orthotopic U87MG-EGFRvIII tumors were treated with resections seven days after tumor inoculation and received either resections alone, resection with intravenous (IV) tail vein delivery of 2M CAR-T cells, resection with local delivery of 2M CAR-T cells in dECM, or resection with local delivery of 200k CAR-T cells in dECM (Fig. 5A). Groups treated with local delivery of CAR T cells contained significantly more responders compared to the peripherally treated group and resection only animals (Fig. 5B-C). Interestingly, animals receiving the low dose of dECM-loaded CAR T cells (200k) had the greatest survival, though both local delivery dose groups had significantly enhanced survival compared to control and peripherally delivered groups (Fig. D).

**Fig. 5.**
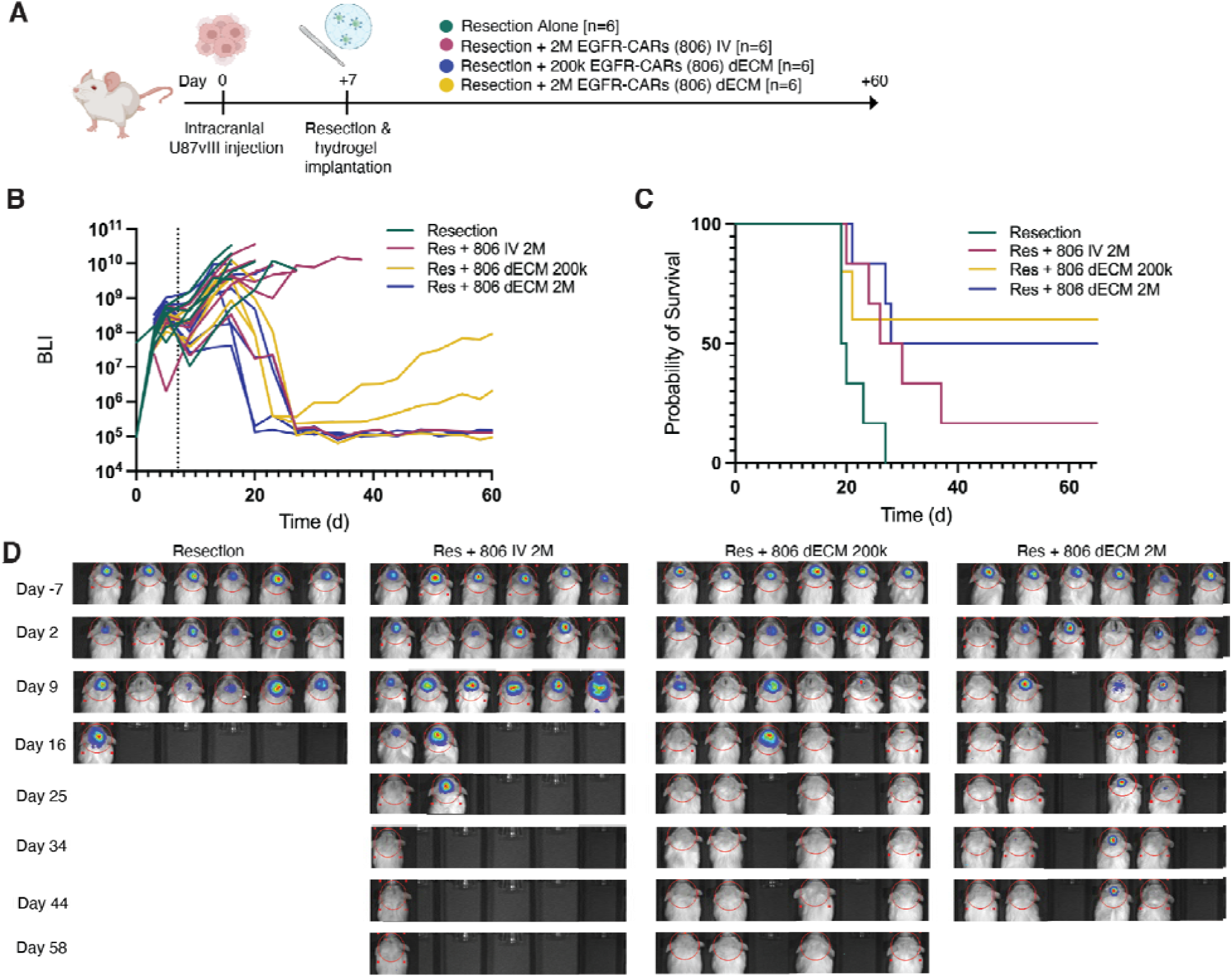
A singular locoregional dose of CAR-T cells in dECM hydrogels controls tumor recurrence and enhances survival outcomes at multiple dose levels. (**A**) Schematic representation of the experimental design. U87vIII cells were transplanted into NSG mice. After tumor growth over 7d, mice were given incomplete surgical resection alone, surgical resection with intravenous 2×10^6^ EGFR-CAR T cells, surgical resection with local dECM hydrogel containing 2×10^5^ EGFR-CAR T cells, or surgical resection with local dECM hydrogel containing 2×10^6^ EGFR-CAR T cells (n=6 per group). Mice were followed with BLI until 60d or until humane endpoints were reached. (**B**) BLI signal representing tumor burden measured in individual mice over time. Vertical lines indicate time of surgical resection and treatment. (**C**) Kaplan-Meier survival curves of animals treated with resection only, resection with high dose IV EGFR-CAR, resection with low dose EGFRgels, and resection with high dose EGFRgels. Data are analyzed with the log-rank Mantel-Cox test. (**D**) Representative images of BLI of all mice over time, red circles highlight tumor location.

### Single dose of locally delivered EGFR CARs prevents recurrence equivalent to repeated peripheral dosing

Local immunotherapeutic tools for GBM have great promise, but access to the intracranial space is limited. Surgical resections cannot be performed repeatedly for multiple local doses. In order to further investigate the tumor control achieved when comparing local and peripheral dosing, a fourth cohort of xenograft mice were treated with a single local dose of CAR T cells (dECM) or multiple peripheral doses (IV). Seven days after tumor inoculation with U87MG-EGFRvIII tumor cells, these mice received either resections alone, resections followed by IV tail vein delivery of 2×10^6^ CAR T cells every 4d, or resections with a one-time local dECM injection of 2×10^6^ CAR T cells (Fig. 6A). Mice were monitored via BLI until study or humane endpoints. Both of the CAR T cell treated groups showed study-long tumor control despite a large difference in total dose (Fig. 8B). One-time local delivery of 2×10^6^ dECM-loaded CAR T cells significantly improved survival outcomes compared to resection only, and was equivalent to mice receiving five peripheral doses of CAR T cell therapy, a total 10×10^6^ cells (Fig. 6C-D).

**Fig. 6.**
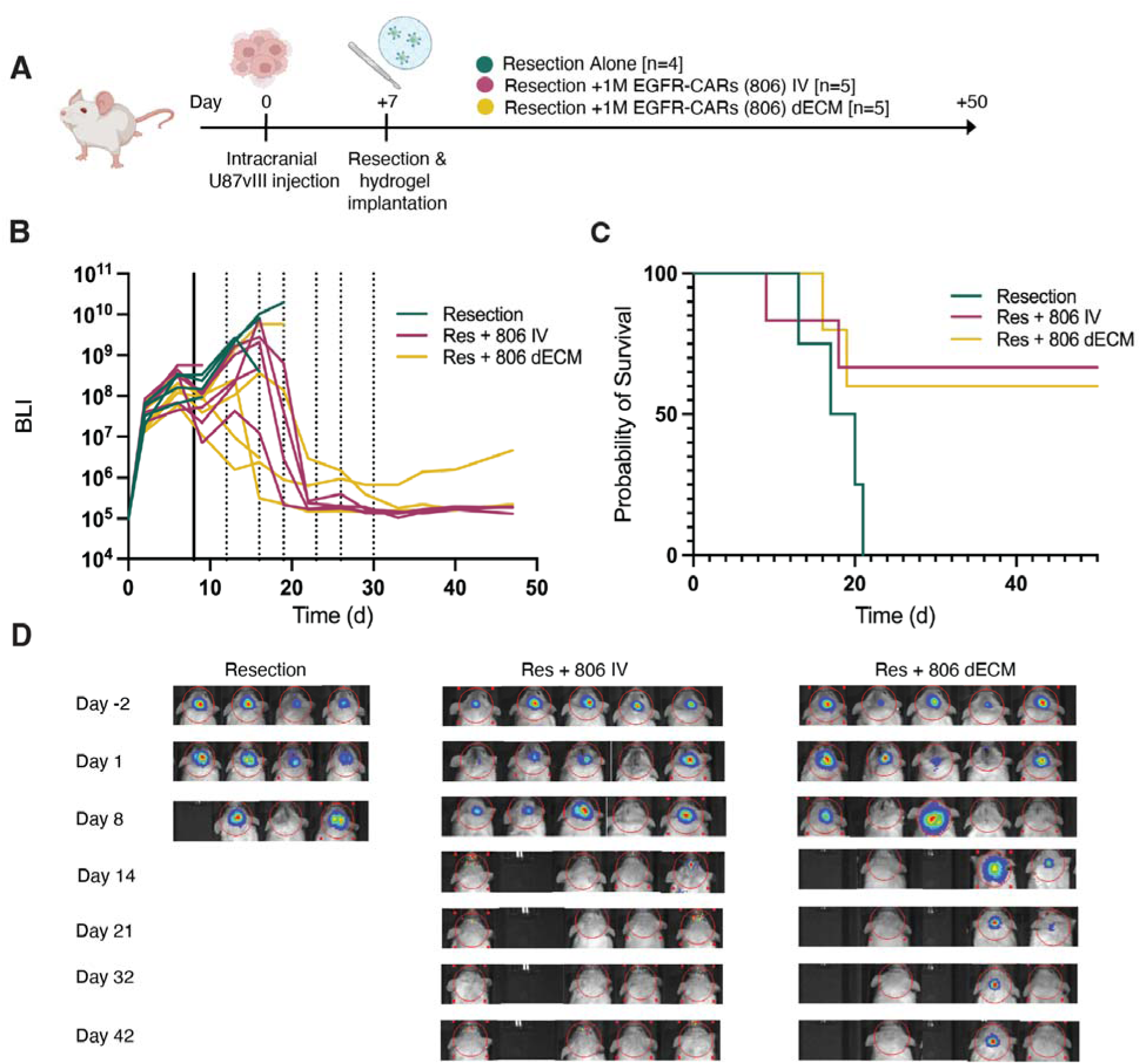
Single locoregional dose of EGFR-CAR-loaded dECM hydrogels after resection confers equivalent survival benefit with repeat IV infusions of EGFR CAR T cells into post-resection GBM xenograft mice. (**A**) Schematic representation of the experimental design. U87vIII cells were transplanted into NSG mice. After tumor growth over 7d, mice underwent incomplete surgical resection alone, surgical resection with repeated intravenous 1×10^6^ EGFR-CAR-T cells, or surgical resection with one-time local dECM hydrogels containing 1×10^6^ EGFR-CAR-T cells (n=5 per group). Mice were followed with BLI until 60d or until humane endpoints were reached. (**B**) BLI signal representing tumor burden measured in individual mice over time. Solid vertical line indicates time of surgical resection and hydrogel placement while dashed vertical lines indicate repeated peripheral infusions for IV-treated mice. (**C**) Kaplan-Meier survival curves of control, resection only, resection with IV EGFR-CARs, or resection with EGFRgel treatment. Data are analyzed with the log-rank Mantel-Cox test. (**D**) Representative images of BLI of all mice over time, red circles highlight tumor location.

Taking all the presented *in vivo* data together, we conclude that CAR T cells delivered locally into GBM resection cavities provide effective and durable tumor control with longer survival at a lower dose than required by peripheral delivery.

## DISCUSSION

City of Hope pioneered the intracranial locoregional delivery (via a Rickham reservoir) of CAR T cells to glioma, and other lead clinical programs are increasingly utilizing locoregional delivery methods to either the ventricles or resection cavity(*7, 27*). Placement of Ommaya or Rickham reservoir systems into the ventricles or resection cavity for locoregional CAR T cell infusion is currently being investigated in multiple clinical trials targeting tumors of the CNS to control local recurrence and allows for repeatable locoregional infusions(*7, 27–31*). Intracavitary therapy using IL13Rα2 CAR T cells led to local tumor control without the adverse events often associated with peripheral infusions, such as the severe cytokine release syndrome (CRS) or neurotoxicity that is seen in patients with high disease burden(*7*). A recent first-in-human clinical trial evaluating feasibility and safety of bivalent CAR T cells against recurrent GBM demonstrated that the intrathecal delivery of CAR T cells is feasible without severe CRS symptoms at multiple dose levels. Additionally, the majority of patients also exhibited robust evidence of bioactivity that is still being evaluated over time(*32*). However, the Rickham or Ommaya devices can be problematic to access subcutaneously, can impair incision healing by increasing the likelihood of wound drainage, and can become a nidus for infection. Similarly, the Gliadel carmustine chemotherapy-loaded wafers can be left within the post-resection space for sustained release into local brain tissue. These wafers elute the majority of chemotherapy payload within seven days, but the remaining empty wafer is not meant to degrade and may increase risk of local adverse events. Therefore, improvements in local delivery strategies are required for the field to exploit all therapeutic opportunities while prioritizing minimizing harm to patients.

To build upon methods for the locoregional delivery of CAR T cells, biomaterial delivery platforms have begun to facilitate methods to improve the local expansion of transplanted cells and enhance cell retention at resection site margins. Almost every patient receives at least one surgical debulking intervention, and intracavitary injections can be easily integrated into current surgical procedures and clinical practice. Despite complete resection of contrast-enhancing tumor, the majority of recurrences are local and within centimeters of the surgical margins, suggesting that more aggressive, cavity-focused treatments may promote progression-free survival(*33, 34*). Although preclinical studies have shown that, based on cell dosing, locoregional delivery is more effective than systemic delivery in treating CNS tumors(*35, 36*), there is still significant uncharted territory regarding intervention timing, efficacy, dosing, and on-target/off-tumor effects that requires rigorous preclinical evaluation.

Novel delivery systems are needed to reduce side effects and improve the accuracy of the immunotherapies that are becoming increasingly popular for the treatment of gliomas. The co-delivery of CAR T cells within a biomaterial vehicle is an innovative approach to locoregional administration and the treatment of solid tumors. Commercially available hydrogel solutions are indispensable to practices such as organoid tissue culture. However, the tumor-derived origin and batch-to-batch variation of products like Matrigel (Corning) limits its use in human studies. Naturally-derived biomaterial-based systems such as dECM can not only provide specific and targeted delivery of cargo, but also extracellular signaling moieties to boost immune response with low toxicity. ECM scaffold proteolysis, degradation, and discharged peptide motifs may increase vascular permeability, induce angiogenesis, and modulate an innate immune response(*37*). Within the resection cavity, ECM scaffolding could allow for increased targeting of residual tumor cells, as well as potentiate the effects of delivered CAR T cells by recruiting antigen-presenting cells and other immune modulators into an otherwise immunosuppressive environment.

Our results demonstrate the significant therapeutic potential of delivering cell therapies locally into the GBM surgical resection cavity. Three mechanisms contribute to cell death during and after transplantation: mechanical forces exerted on free cells during injection, anchorage-dependent cell death, and lack of support from the degenerative host tissues(*38–40*). Peripherally delivered CAR T cells have multiple barriers to enter the brain, let alone penetrate the tumor parenchyma. Local delivery removes those challenges, leaving only the tumor microenvironment as the main opposition to treatment response. Previous studies have demonstrated that the direct transplantation of ECM-derived hydrogel carrying cells into CNS lesions significantly improves cell survival, compared to cells delivered without a supportive scaffold.

Here we show that locally delivered EGFR-CAR T cells can create durable anti-tumor responses in our orthotopic GBM mouse models, above and beyond the responses that we see in animals treated with IV CAR T cells. Mice treated with EGFR-CAR T cells in a dECM hydrogel were the only treatment group CD3+ T cells present within tumor-bearing tissues at endpoint, compared to animals treated with EGFR-CAR T cells through peripheral IV tail vein administration. When the locally administered dosage of cells is reduced 10-fold, a similarly durable or even improved response is seen as compared to the higher single or repeated doses delivered intravenously. With an intracavitary treatment delivered conveniently at time of resection that mimics natural brain architecture, there is an early chance to address residual tumor using a lower dose of targeted cells in a vehicle that boosts immune cell recruitment and function while preventing off-target toxicities and systemic effects.

This study does contain limitations that must be considered when interpreting our data. First, the NSG mice models do not adequately mimic the full complement of the human immune system and thus these studies alone will not fully depict the immune cell interactions and the GBM tumor microenvironment. However, patients with brain injuries, including tumors such as GBM, are often in a state of relative immune dysfunction that can be perpetuated with chemotherapies and radiation, making it difficult to accurately model GBM, Second, the path of development for a delivery device requires significant evaluation of long-term safety and potential toxicity of biogels delivered to the brain. Nonetheless, these data illustrate potential opportunities to focus research and development when developing immunotherapy delivery vehicles. Importantly, there is tunability to the dECM system. The biogel formula could be expanded upon to potentially include the addition of a matrix or soluble protein components that would enhance T cell fitness and duration of response. The ECM consistently undergoes remodeling to maintain tissue homeostasis, and impaired ECM remodeling is a hallmark of disease and cancer. Most immune cells express a diverse comportment of ECM receptors in order to regulate cell movement and effector functions, and regulation of these functions by ECM proteins is evidenced by our NK cells performing sub-optimally when encapsulated within collagen-only hydrogels, as increased collagen deposition has been linked to decreased NK immunosurveillance(*41, 42*).

In summary, locoregional delivery of CAR T cells can be safely and effectively included within routine surgical interventions for GBM to improve tumor response after resection. Maximally safe surgical resections are the goal of GBM SOC treatment, but the extent of resection is often limited by eloquent areas and the diffusely infiltrative nature of gliomas. In the complex equation that surgeons must balance between disease control and patient neurological outcome, this adjuvant treatment could buttress resection surgery and bridge the gap to the administration of systemic therapies or radiation. Compared to other methods of locoregional delivery for immune- and chemotherapies, we have demonstrated that our treatment has the power to expand, seek, and destroy remaining tumor cells with improved persistence. Treating the surgical resection cavity with CAR T cell-loaded hydrogel scaffolds could allow for enhancement of survival by providing the beneficial therapeutic effects of immunotherapy delivered at a convenient time with the fewest number residual cells before the tumor can adapt and regrow.

## MATERIALS AND METHODS

### Study Design

In this study, we developed an intracavitary delivery vehicle for local administration of CAR T cells into GBM resection cavities. We characterized the biochemical and structural components of our hydrogel delivery vehicle, and then the release, viability and potency of cells after encapsulation within the delivery vehicle. Finally, we evaluated the antitumor potency of this delivery strategy compared to the clinical standard in postoperative GBM. For all *in vitro* assays, three to six replicates for each condition were used to evaluate intra group variability and allow for defining statistical significance. For all *in vivo* experiments, five to six mice were used in each treatment group to ensure statistical power when evaluating tumor recurrence and survival probability.

### Hydrogel Fabrication

Native fresh frozen pig brain tissue was purchased through XX. Decellularization reagents including sodium dodecyl sulfate (SDS), porcine pepsin, and hydrochloric acid (HCl) were purchased from Sigma Aldrich. Frozen native tissues were stored in a freezer at −80°C for at least 24h before beginning the decellularization process. Once thawed, porcine brain tissue was dissected into smaller portions of approximately 1 cm^3^ and then rinsed in 500 mL of 1x PBS three times to remove any gross debris. Brain tissues were then moved into a decellularization solution containing 0.1% SDS (Sigma Aldrich) and 1% pen-strep (Corning) in 1x PBS. The solution containing tissues was placed onto an orbital shaker at 80 RPM. Supernatant containing cellular debris was decanted and replaced every 24h for 4d. After decanting a final time, remaining tissue was washed three times in 1x PBS and centrifuged at 4000 RPM for 5 min. Conical tubes were decanted and refilled with deionized water, then centrifuged again. These washing steps were repeated 10 times to remove residual SDS from the tissues. Final decanted tissues were aliquoted into several conical tubes in deionized water and frozen at −80°C for at least 24h. Frozen tissues were lyophilized for 48h and stored dry at −20°C until use.

To prepare pre-gel solutions, lyophilized and decellularized tissue must be solubilized into protein monomeric components. Lyophilized tissue was powdered using a small handheld grinder, and then dissolved as 10:1 weight ratio into 1mg/mL pepsin solution (Sigma) within 0.01M HCl for 48-96h with stirring. Pepsin digestion was complete when the liquid solution is homogeneous with no large visible particles. When digestion was done, the reaction was stopped by raising the pH and salt concentrations to physiologic levels. The resulting pre-gel solution was kept at 4°C until use and will form a gel when the liquid is brought up to 37°C as a result of the collagen kinetics.

### Cell Culture and Cell Encapsulation

The U87MG cell line was obtained from American Type Culture Collection (ATCC HTB-14) and cultured in MEM containing HEPES, sodium pyruvate, GlutaMAX-1, penicillin/streptomycin, and supplemented with 10% fetal bovine serum (FBS) (all purchased from Corning). This cell line was transfected to express the EGFRvIII protein, click beetle green luciferase, and green fluorescent protein through single-cell purification and expansion. Cells were routinely screened for identity and for mycoplasma contamination.

The EGFR-targeting scFv 806 was synthesized and ligated into a pTRPE lentiviral vector with the EF1alpha promoter. The control CAR-CD19 was a gift from the Carl H. June lab at the University of Pennsylvania. All CAR molecules have a second-generation CAR design containing the 4-1BB costimulatory domain with a CD3 zeta chain. Lentivirus was packaged in HEK293T cells using a split genome approach and titered using SUPT1 cells from ATCC (CRL-1942). Healthy human T cells were isolated from PBMCs from the Human Immunology Core at the University of Pennsylvania and were transduced using lentiviral vectors. T cells were stimulated for 5 days using Dynabead Human T-Activator CD3/CD28 (Life Technologies) at a 3:1 bead-to-cell ratio. Cell concentration was determined using a Coulter Multisizer (Beckman Coulter) and maintained at 0.7e6 cells per mL until fully rested in around 300 fL in volume. Cells were cultured in R10 media (RPMI-1640 supplemented with GlutaMAX-1, HEPES, sodium pyruvate, penicillin/streptomycin, and 10% FBS) with 30 IU/mL rhIL-2 (Thermo Fisher Scientific). CAR T cells were then cryopreserved in 90% FBS with 10% DMSO for future use.

### Rheological Characterization

Hydrogels of each treatment were prepared in 1x PBS and kept on ice until use for rheological testing. 500 μL of hydrogel were pipetted onto the testing platform of an Anton Paar Gemini rheometer and warmed to 37°C for gelation before analysis for storage and loss shear moduli using parallel plate geometry. Time sweep and frequency sweep experiments over 0.1 to 100 Hz were performed at 5% strain.

### DNA, Protein Content and Cytokine Measurement

DNA extraction from native and decellularized brain samples was performed following manufacturer’s protocol (KIT). Double-stranded DNA was quantified using the Qiagen DNEasy Blood & Tissue Kit as per kit instructions.

All tissue samples were analyzed for measurement of dominant proteins. 50 mg of lyophilized native and decellularized tissues were suspended in 1x PBS as per kit instructions. Supernatants were collected and buffer was added. Content of proteins was determined via Collagen Assay Kit (Sigma-Aldrich) and Total Glycosaminoglycans Assay Kit (abcam) respectively, following the individual manufacturers’ protocols.

Supernatant from CAR-T cells incubated in hydrogels was analyzed using a MACSPlex Cytokine 12 Kit (Miltenyi Biotec) for released cytokines in culture. Supernatants were incubated with MACSPlex Capture Beads to bind cytokines for 2h, and then incubated with MACSPlex Cytokine Detection Reagents containing APC-conjugated antibodies for 1h for the respective analytes to form sandwich complexes. Samples were analyzed on APC fluorescence for each capture bead population and are compared to standard curves for each cytokine to calculate concentration of cytokines in each sample.

### Scanning Electron Microscopy Imaging

Morphology of gold sputter-coated lyophilized hydrogel samples was examined using a ThermoFisher FEI Quanta 250 FEG Scanning Electron Microscope. Hydrogels were frozen overnight at 80°C and lyophilized. Lyophilized gels were mounted into 10 mm SEM stubs and sputter coated with gold for 15 seconds then imaged under an accelerated voltage of 5 kV. Images were acquired at 500X and 1000X to observe porosity and structure of the hydrogels.

### CAR T Cell Release and Tumor Killing Axion Cell Impedance Assays

Cell release was tested via Transwell assays (Corning). Non-encapsulated and hydrogel encapsulated CAR T or NK cells were incubated in R10 media solution within an insert containing a permeable PTFE membrane with 8.0 μm pore size and allowed to freely diffuse into R10 media containing IL-2 in bottom chamber over time. Released cells were quantified after 48h and 96h. The released cells collected at the 48h time point were checked for viability using Trypan Blue exclusion (Thermo Fisher Scientific) before use in cell killing assays.

U87MG target tumor cells were seeded at 50k/well into Axion Biosystems microelectrode-containing 96-well plates (Axion Biosystems). Each impedance plate was prepared prior to experimentation by coating first with poly-D-lysine for 6h and then coated with 20 μg/mL laminin overnight at 37°C. After coating, wells were rinsed with diH2O and overlaid with 100 μL of cell culture medium for background impedance recording in the Axion Biosystems ZHT analyzer (Axion Biosystems). After a baseline was established, the plate was removed from the analyzer and seeded with 50k target cells in a volume of 100 μL per well for a final volume of 200 μL per well. After cell plating, the plate was left for 1h at room temperature to improve attachment to the bottom surface before being returned to the analyzer for data collection. Data was collected every 1 min for 24h to monitor target cell attachment and monolayer formation. For cytotoxicity co-culture assays, the instrument was paused at 24h and medium was exchanged for medium containing 1:1 dosages of effector (CAR T or NK cells) or non transduced (UTD) control T cells, or medium alone. Changes in impedance were recorded as the resistive component of the complex impedance calculation, as described previously. Using AxIS Z software (Axion Biosystems), all data were corrected for “media alone” control and then normalized to the impedance at the time of addition of effector cells. Percent cytolysis was calculated using the no treatment control and full lysis controls to determine the percent of target cell death.

### In Vivo Experimental Modeling

All mice experiments were conducted according to Institutional Animal Care and Use Committee (IACUC)-approved protocols. For orthotopic glioma modeling, 5e5 U87MG-CBG-GFP cells were implanted intracranially into 8-week-old NSG mice. Intracranial surgical implants were done using a stereotaxic setup with tumor cells implanted into the brain at 2 mm right of and 2 mm anterior to lambda at a depth of 2 mm. Tumor progression was evaluated by luminescence emission on a Xenogen IVIS spectrum after intraperitoneal injection of D-luciferin (Gold Biotechnology) according to manufacturer’s instructions. T cells were administered in a total volume of 10 μL of gel for intracranial treatment or in a volume of 100 μL of PBS via the tail vein for intravenous treatment. Survival was followed over time until the predetermined IACUC-approved endpoint was reached: 50-60d after tumor inoculation or once humane endpoints were reached, whichever happened first.

### Histology and Immunohistochemistry

Brains were explanted from animal studies and preserved in 4% paraformaldehyde at 4C for at least 24h before being transferred to the Molecular Pathology and Imaging Core Facility (University of Pennsylvania) for paraffin embedding, sectioning, and H&E staining. For immunohistochemistry, dehydrated brain sections were rehydrated with two xylene solution changes and then gradually rehydrated in ethanol solution changes (100%, 100%, 90%, 90%, 50%, 50%) for 5 minutes per wash. After that, tissue sections were transferred to deionized water and then placed into antigen retrieval solution for 20 min at 60°C. After antigen retrieval, slides were washed in cool deionized water and then prepared for blocking.

For immunofluorescence staining, tissue sections were washed with TBS containing 0.1% Tween-20 (v/v) then permeabilized and blocked using a solution containing 10% donkey serum (v/v), 0.5% Triton X-100 (v/v), 1% BSA (w/v), 0.1% gelatin (w/v), and 22.52 mg/mL glycine in TBST for 1h at room temperature. Tissue sections were incubated with anti-CD3-APC antibody (Miltenyi Biotec) diluted in TBST with 5% donkey serum (v/v) and 0.1% Triton-X100 (v/v) overnight at 4°C. After washing with TBST, tissue sections were DAPI stained and then washed and mounted in Fluoromount-G Mounting Medium (Thermo Fisher Scientific), coverslipped, and sealed with nail polish for imaging. Images were acquired on a Zeiss LSM 710 confocal microscope and analysis was performed using ImageJ Fiji 2.9.0.

### Statistical Methods

All results are presented as means ± SEM unless denoted otherwise. Error bars represent the SD of the mean from independent samples and animals are randomly grouped prior to the initiation of treatment. Two-tailed Student’s t tests were used to test statistical significance of differences between two groups, with Tukey’s post hoc tests. One-way analysis of variance (ANOVA) was performed for multiple comparisons of three treatment groups or more. Log-rank (Mantel-Cox) tests were used to analyze statistical significance across groups for survival analysis. All statistical analyses were performed in Prism (Prism 10.2.2, GraphPad Software). Statistical significance was determined at * p<0.05, ** p<0.01, *** p<0.001, **** p<0.0001.

## List of Supplementary Materials

Fig. S1. Decellularization procedures to create dECM hydrogels.

Fig. S2. EGFR CAR construct and flow cytometry validation of CAR expression.

Fig. S3. Resection of GBM alone does not enhance survival outcomes in intracranial xenograft mice.

Fig. S4. dECM hydrogels injected into the flank and brain demonstrate a lack of long-term toxic and local inflammatory effects.

## Acknowledgments

The authors would like to thank the Flow Cytometry Core Facility, the Cellular and Development Microscopy Core Facility and the Stem Cell and Xenograft Core Facility at the University of Pennsylvania for their support and equipment contributions to this work.

## Funding

Breakthrough Challenge Award (ML)

Abramson Cancer Center Glioblastoma Translational Center of Excellence (MTL, SLB, KH, JP, LZ, DMO, ZAB.)

Templeton Family Initiative in Neuro-Oncology (DMO)

Maria and Gabriele Troiano Brain Cancer Immunotherapy Fund (DMO)

National Institutes of Health grant R37CA285434 (ZAB)

## Author contributions

Conceptualization: MTL, ZAB, DMO

Methodology: MTL, ZAB, DMO

Experimentation: MTL, SLB, KH, JP, LZ

Funding acquisition: MTL, ZAB, DMO

Supervision: DMO, ZAB

Writing and Editing: MTL, SLB, ZAB, DMO

## Data and materials availability

All data are available in the main text or the supplementary materials.

