## Supplemental Figures for "CAR-T cells locally delivered in porcine decellularized matrix hydrogels enhance survival in post-resection glioblastoma"

**List of Supplementary Materials**

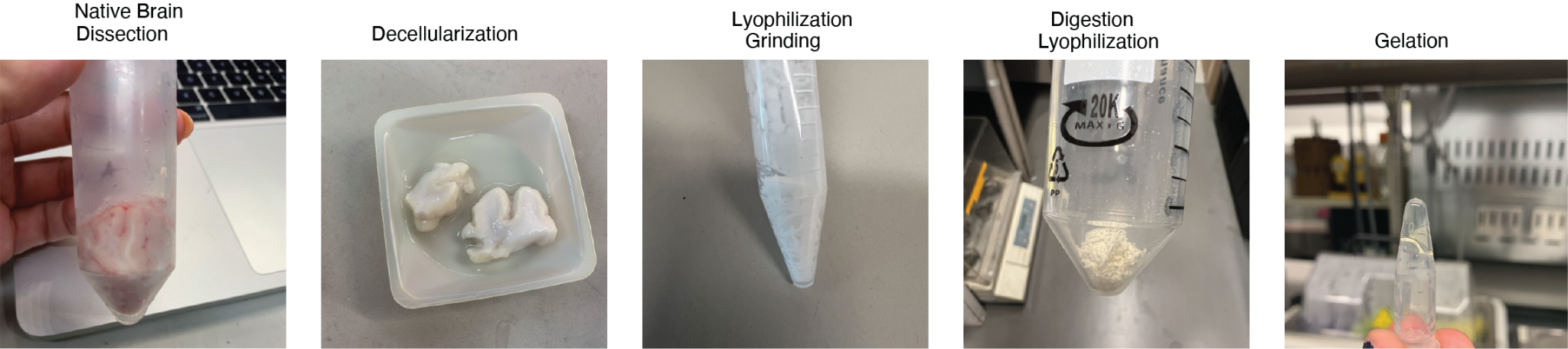

**Fig. S1. Summary of decellularization procedures to create dECM hydrogels.**

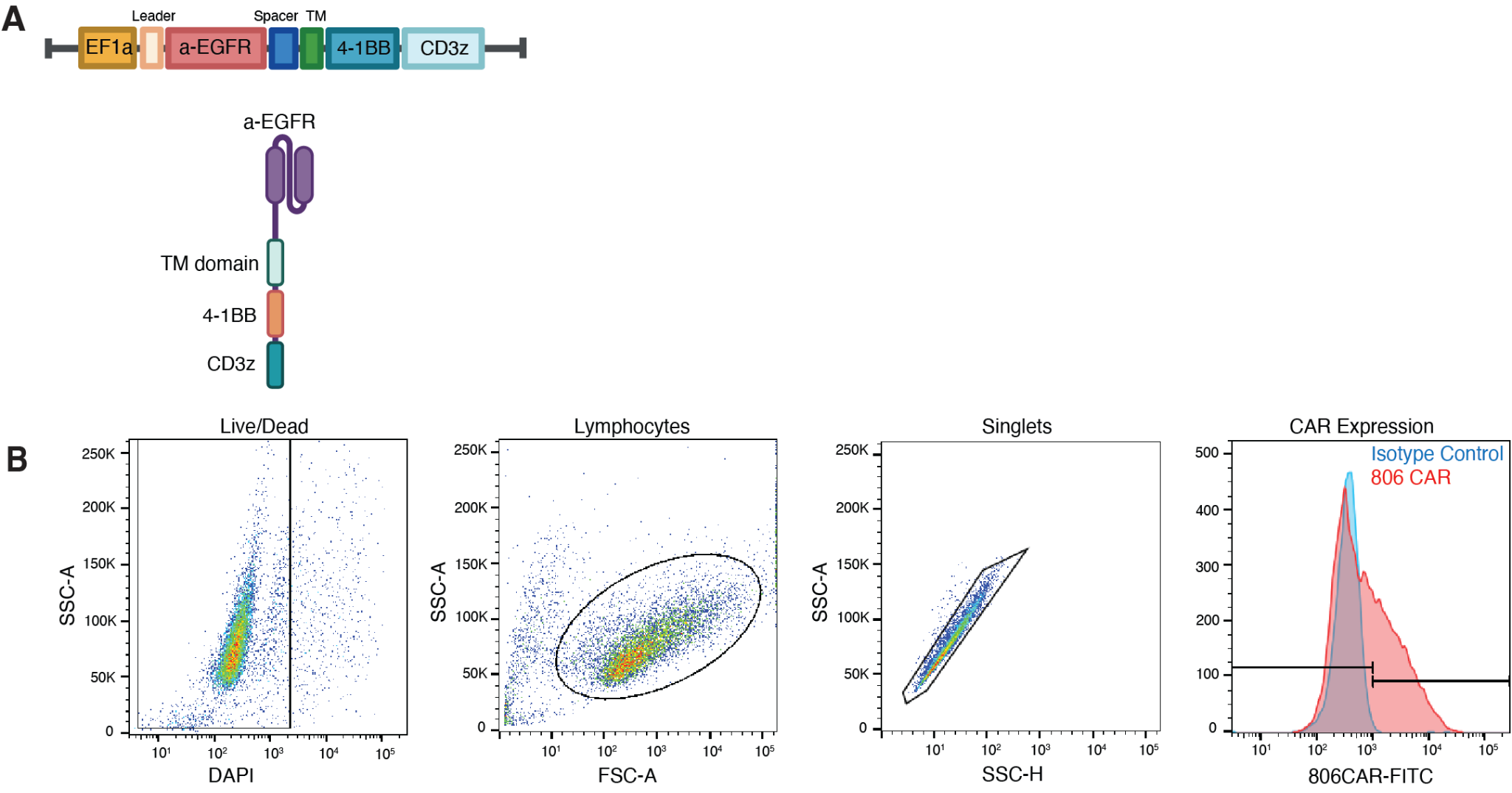

**Fig. S2. EGFR CAR construct and flow cytometry validation of CAR expression.** (A) Diagram of CAR plasmid and subsequent CAR construct. (B) Quantitative analysis of EGFR CAR density in T cells after transduction protocols using flow cytometry.

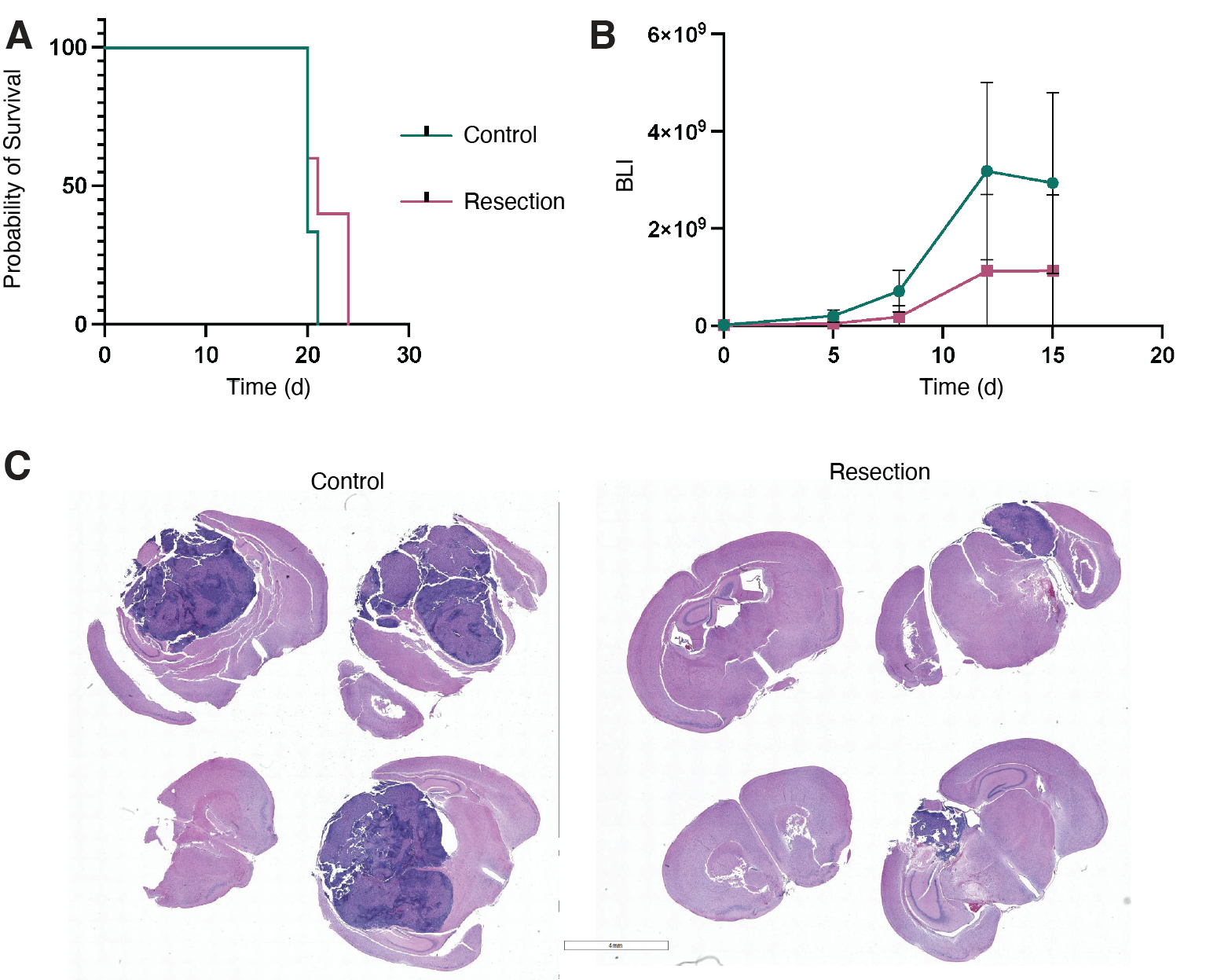

**Fig. S3. Resection of GBM alone does not enhance survival outcomes in intracranial xenograft mice.** (A) Kaplan-Meier survival curves of tumor-bearing mice either receiving no treatment or only surgical resection treatment. (B) BLI signal intensity in mice after receiving either no treatment or only surgical resection procedure. Data are shown in means ± SD (n=5). (C) H&E-stained images of brain tissues from U87MG-bearing mice without (left) and with (right) surgical resection, taken at humane endpoint. Scale bar 4mm.

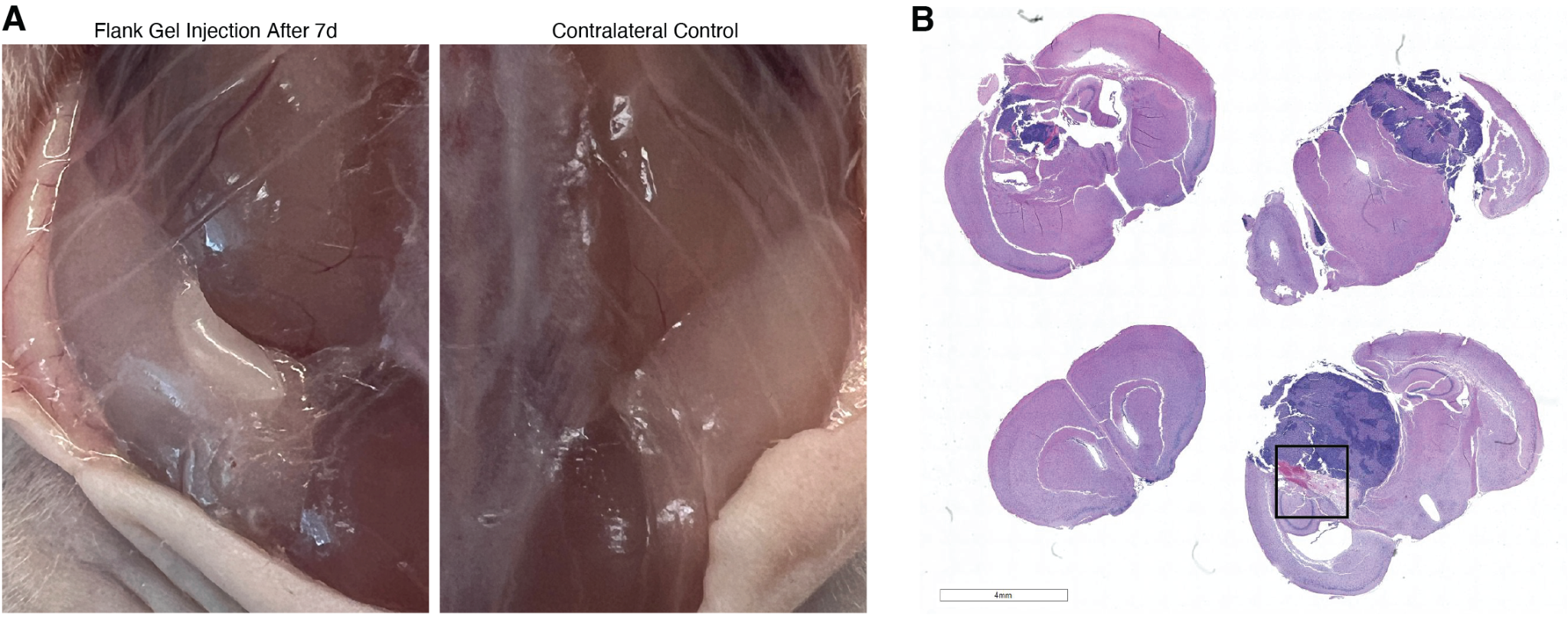

**Fig. S4. dECM hydrogels injected into the flank and brain demonstrate a lack of long-term toxic and local inflammatory effects.** (A) Flank injections of empty dECM hydrogels demonstrate lack of adverse reaction to mice after 7d compared to contralateral control areas. (B) H&E staining of tumor-bearing brains isolated after treatment with surgical resection and empty dECM hydrogels. Scale bar 4mm. Some hydrogel presence remains at 21d (boxed).
